# ECMarker: Interpretable machine learning model identifies gene expression biomarkers predicting clinical outcomes and reveals molecular mechanisms of human disease in early stages

**DOI:** 10.1101/825414

**Authors:** Ting Jin, Nam D. Nguyen, Flaminia Talos, Daifeng Wang

**Affiliations:** Department of Biostatistics and Medical Informatics, University of Wisconsin – Madison, Madison, WI, 53706, USA; Department of Computer Science, Stony Brook University, Stony Brook, NY, 11794, USA; Departments of Pathology and Urology, Stony Brook Medicine, Stony Brook, NY, 11794, USA; Stony Brook Cancer Center, Stony Brook Medicine, Stony Brook, NY, 11794, USA; Waisman Center, University of Wisconsin – Madison, Madison, WI, 53705, USA

**Keywords:** Interpretable machine learning, early cancer, gene expression biomarkers, gene regulatory network, semi-restricted boltzmann machine, cancer medicine

## Abstract

Gene expression and regulation, a key molecular mechanism driving human disease development, remains elusive, especially at early stages. Integrating the increasing amount of population-level genomic data and understanding gene regulatory mechanisms in disease development are still challenging. Machine learning has emerged to solve this, but many machine learning methods were typically limited to building an accurate prediction model as a “black box”, barely providing biological and clinical interpretability from the box. To address these challenges, we developed an interpretable and scalable machine learning model, ECMarker, to predict gene expression biomarkers for disease phenotypes and simultaneously reveal underlying regulatory mechanisms. Particularly, ECMarker is built on the integration of semi- and discriminative- restricted Boltzmann machines, a neural network model for classification allowing lateral connections at the input gene layer. This interpretable model is scalable without needing any prior feature selection and enables directly modeling and prioritizing genes and revealing potential gene networks (from lateral connections) for the phenotypes. With application to the gene expression data of non-small cell lung cancer (NSCLC) patients, we found that ECMarker not only achieved a relatively high accuracy for predicting cancer stages but also identified the biomarker genes and gene networks implying the regulatory mechanisms in the lung cancer development. Additionally, ECMarker demonstrates clinical interpretability as its prioritized biomarker genes can predict survival rates of early lung cancer patients (*p*-value < 0.005). Finally, we identified a number of drugs currently in clinical use for late stages or other cancers with effects on these early lung cancer biomarkers, suggesting potential novel candidates on early cancer medicine. ECMarker is open source as a general-purpose tool at https://github.com/daifengwanglab/ECMarker.

## 1 Introduction

Human disease development such as cancer is a complex, dynamic process that is fundamentally driven by abnormal molecular mechanisms. However, understanding the cancer mechanisms is still a challenging task, especially during the early cancer development (Koeffler, McCormick and Denny 1991, Herbst, Morgensztern and Boshoff 2018). To this end, the tumor/node/metastasis (TNM) system has been widely used to characterize and classify the cancer development into various stages (Ludwig and Weinstein 2005). The TNM stages were further associated with a number of individual molecular biomarkers and clinical outcomes such as survival rates (Ludwig and Weinstein 2005). However, with the advancement of whole genome sequencing of numerous human tumors, it became apparent that the molecular profiles of various tumor stages might not necessarily be reflected by the TNM system. Therefore, using systems biology approaches to identify the biomarkers that drive cancers from early to late stages could allow for better understanding of cancer mechanisms and offer new venues for the development of new preventive and therapeutic strategies.

However, there is a gap in understanding of the molecular biomarkers of early cancer and their underlying mechanisms at a system level. For example, the lung cancer, causing 27% of cancer-related deaths in the USA alone (Siegel, Miller and Jemal 2018), is localized to the lung; neither lymph nodes nor other organs are believed to be affected at the early stage. As the cancer progresses to a more advanced stage, nearby lymph nodes and other organs may be affected (Hu et al. 2008, Frost et al. 1984). This pathological difference suggests that the underlying molecular mechanisms of the early and late stages are different. Also, if cancer is diagnosed at an early stage, such as a localized stage, the five-year survival rate is approximately 50%; this is mostly due to surgical interventions involving lung removal. After the localized stages, survival rates decrease rapidly as cases involving the lymph nodes or other metastatic sites necessitate elaborate treatment strategies (Hu et al. 2008). Nearly 70% of patients with lung cancer present with locally advanced or metastatic disease at the time of diagnosis (Molina et al. 2008). Thus, although a number of studies have indicated that early localized stages are easier to treat and have better survival rates, the underlying molecular mechanisms remain elusive. Here, we hypothesized that early cancers have different molecular wiring at a system level and that understanding this wiring could reveal new biomarkers and mechanisms of early cancer development. Thus, it is essential to identify the specific biomarkers of early cancer to understand the molecular mechanisms driving cancer development; this would enhance early cancer diagnosis and therefore improve survival rates.

Detecting early cancer biomarkers, however, involves the inherent challenges of relating the complex, multi-dimensional molecular processing that occurs in organs and tissues during early-stage cancer to observable clinical phenotypes in human patients. In particular, differential, temporal, and spatial gene expression during early cancer result from disruptions in the complex, dynamic, and multi-gene process that tightly regulates and controls the developmental integrity of organs and tissues. These temporal and spatial gene expression dynamics are fundamentally controlled by a variety of molecules called gene regulatory factors, including transcription factors (TFs) and non-coding RNAs. These factors cooperate in a gene regulatory network (GRN) to carry out correct developmental functions on a genome scale (Iyer, Osmanbeyoglu and Leslie 2017). The nodes of a GRN are genes, and the edges of a GRN connect regulatory factors to their target genes. Disruption of the cooperation between genes and regulatory factors in a GRN can give rise to abnormal gene expression, such as that which is present in diseases such as cancer. Therefore, a fundamental challenge for uncovering early cancer mechanisms is that of understanding the gene regulatory mechanisms, especially GRNs, controlling the changes in gene expression across cancer stages.

The collection of next-generation sequencing (NGS) data from large cohorts such as TCGA (Liu et al. 2018b) provides measurements across multi-omics, including transcriptomics and epigenomics. This allows for studies of temporal dynamics in gene expression and regulation during cancer development and also for the systematic identification of stage-specific cancer biomarkers. Progress has been made in identification of some stage-specific molecular biomarkers of lung cancer, but systematic genome-wide analyses for identification of all potential early-stage biomarkers with predictive value for disease outcome are limited. For example, dysregulations in the epidermal growth factor receptor EGFR, associated with sensitivity of lung cancers to the tyrosine kinase inhibitor gefitinib (Iressa) (Pao et al. 2004), echinoderm microtubule-associated protein-like 4 (EML4), and anaplastic lymphoma kinase (ALK) are frequently involved in oncogenic transformation (Lindeman et al. 2013). Additionally, v-raf murine sarcoma viral oncogene homologue B1 (BRAF) is a driver mutation gene in lung adenocarcinoma (Paik et al. 2011). Although a challenging task, finding novel ways to integrate the large-scale data provided by human tumors would enable the discovery of genome-wide early cancer biomarkers and underlying GRNs.

Traditionally, correlation-based models have been used to select biomarker genes involved in cancer development; e.g., 62 genes were uncovered in this way to distinguish between the early and late stages of clear cell renal cell carcinoma (ccRCC) (Rahimi and Gönen 2018) (Jagga and Gupta 2014). However, correlation-based models only reveal linear relationships, whereas cancer development is a complex, nonlinear process. Thus, machine learning has emerged as a powerful tool to predict biomarkers for various cancer features related to clinical presentation and staging; this tool has been found to be of great help in the diagnosis and treatment of various diseases (Libbrecht and Noble 2015). For example, (Statnikov, Wang and Aliferis 2008) applied random forests (RFs) and support vector machines (SVMs) to microarray data in order to aid in cancer diagnosis. (Xiao et al. 2018) constructed a multi-model ensemble approach to predict cancer in both normal conditions and tumor conditions. However, none of these studies revealed novel cancer mechanistic insights; these studies were limited to building an accurate classification model as a “black box” but lacked any biological or clinical interpretability from the box. In addition, the biological datasets especially for genomics have the challenging of “curse of dimensionality” (Clarke et al. 2008); e.g., variables (e.g., genes) are much more than samples. To solve this, many machine learning methods applied prior feature selections to reduce the dimensionality, which however likely miss potentially important information at the system level.

To address these challenges, we designed a novel, interpretable machine learning approach, ECMarker, that can be used to discover gene expression biomarkers for the early disease stages, and simultaneously unravel the underlying molecular mechanisms in the “black box” such as gene regulatory networks (GRNs). In particular, ECMarker is built on a neural network model, semi-restricted Boltzmann machine (SRBM) allowing lateral connections at the input gene layer for classifying disease phenotypes using population-level gene expression data. The SRBM model (Osindero and Hinton 2007) has been used in non-biological contexts (e.g., computer vision, image classification) enabling modeling intra-connections among input variables (e.g., image patches). Based on the neural network connectivity, ECMarker further enables prioritizing genes and revealing the underlying gene network (from lateral connections) for predicting phenotypes. Compared to other methods, ECMarker is an interpretable model aiming to reveal underlying molecular mechanisms (e.g., gene regulation) while predicting phenotype. Many existing machine learning models still aim to learn a “black box” with high accuracy, which is not straightforward to provide any biological insights as described above. ECMarker, instead, was designed to achieve all “interpretability”, “accuracy” and “scalability” via (1) using the lateral connections at the visible layer (i.e., genes) to reveal gene networks, (2) simultaneously trying to achieve relatively high accuracy of classifying disease stages and (3) inputting all genes and prioritizing genes by implicit feature selection. With applications to cancer genomic data, the prioritized genes and networks for early/late cancer stages revealed potential cancer stage-specific gene biomarkers and GRNs. Furthermore, we found the drugs that have significant effects on the ECMarker biomarkers for uncovering novel genomic medicine to early cancer development.

## 2 Methods and Materials

### 2.1 ECMarker, an interpretable machine learning model to identify gene expression biomarkers and underlying molecular mechanisms for disease development and outcomes

ECMarker consists of three major components (Fig. 1) including (1) a neural network model, integrating the semi-restricted Boltzmann machine (SRBM) (Osindero and Hinton 2007) with the Discriminative restricted Boltzmann machine (DRBM) (Larochelle and Bengio 2008) for classifying disease phenotypes from the gene expression data at the population level; (2) the prioritization of gene expression biomarkers for each phenotype using the integrated gradient method based on the neural network connectivity, and identification of a gene network using the lateral connections at the input layer; (3) the functional and survival analyses of biomarker genes and networks for revealing underlying molecular mechanisms in the disease phenotypes (biological interpretability) and predicting clinical outcomes (clinical interpretability). We elaborated each component as follows.

**Fig. 1.**
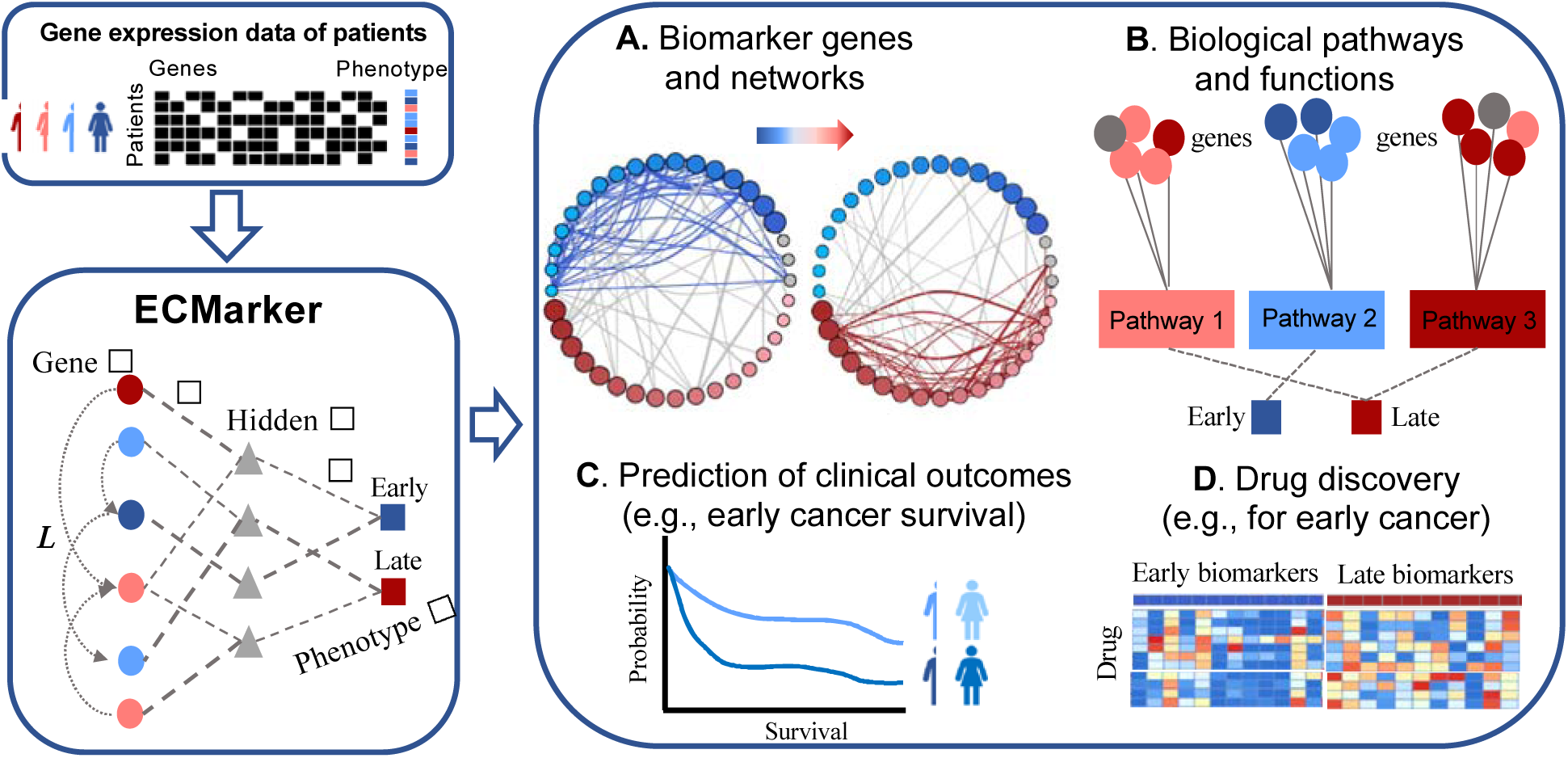
ECMarker, an interpretable machine learning framework for the identification of gene expression biomarkers of cancer stages and the prediction of clinical outcomes. ECMarker is a hierarchical neural network approach integrating semi- and discriminative-restricted Boltzmann machine models, to input the gene expression data of patients for predicting their disease phenotypes; e.g., early and late cancer stages. In particular, the ECMarker classification model consists of three layers: (1) the input gene layer ***v***, (2) the hidden layer ***h***, and (3) the output phenotype layer ***y***; e.g., early vs. late cancer stages. The lateral connections at the input gene layer enable identifying a gene network providing potential mechanistic insights for disease phenotypes. Thus, in addition to the phenotype prediction, ECMarker is also biologically and clinically interpretable for (**A**) identifying the gene expression biomarkers and gene networks for phenotypes (e.g., early and late stages); (**B**) revealing the associated biological functions and pathways for each phenotype; (**C**) predicting clinical outcomes such as survival rates, especially for early cancer patients; and (**D**) discovering novel drugs potentially affecting early cancer. Red and blue represent early and late stages, respectively.

### 2.2 The ECMarker classification model

The standard restricted Boltzmann machine (RBM) is an energy-based model that uses a layer of *n* hidden units to model a distribution over *m* visible units in the other layer (Hinton and Salakhutdinov 2006). The connections between the two layers (i.e., visible layer to hidden layer) and all visible and hidden units form a bipartite graph. Thus, the connections within a layer (e.g., a visible unit to another visible unit) are prohibited. Also, the standard RBM typically takes binary values; i.e., ***v*** ∈ {0,1}^*m*^ and ***h*** ∈ {0,1}^*n*^, and is often trained by the input distributions only. In the ECMarker, we have extended the RBM, based on the semi-restricted Boltzmann machine (SRBM) (Osindero and Hinton 2007) and the Discriminative RBM (DRBM) (Larochelle and Bengio 2008) for enabling (1) classification, (2) inputting continuous values of visible units (e.g., gene expression) and (3) modeling the gene relationships (i.e., a network) as follows. First, ECMarker inputs the expression profiles of *m* genes as *m* visible nodes ***v*** ∈ ℝ^*m*^. Second, the hidden layer in the ECMarker consists of the binary variables ***h*** ∈ {0,1}^*n*^, where *n* is number of hidden nodes. Finally, the output layer ***y***∈ {0,1}^*K*^ consists of all *K* phenotypes to predict. The improvements of the ECMarker classification model include:

1. To deal with the real-valued gene expression data, we replaced the binary visible units in the RBM by linear units with independent Gaussian noises. To simplify calculation, we used the Gaussian noise ∼𝒩(0,1) and transformed the input data before training; i.e., standardizing features by removing the mean and scaling to unit variance.
2. We added an output layer ***y***∈ {0,1}^*K*^ with discretized values modeling *K* phenotypes (e.g., *K*=2, early vs. late disease stages) on the top of the hidden layer, and then used a joint distribution 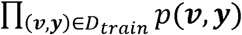 over the training data set *D*_train_ of the input ***v*** ∈ ℝ^*m*^ (e.g., *m* gene expression values) and associated phenotype ***y*** for classification.
3. We allowed lateral connections among the visible units as SRBM did for modeling a network linking genes while predicting phenotypes.

In particular, the probability distribution represented by the ECMarker classification model with parameters *Θ* is *p*(***v, y, h***|*Θ*) ∝*e*^-*E*(***v,y,h***;*Θ*)^, where *E*(***v, y, h***; *Θ*) is the energy function defined by

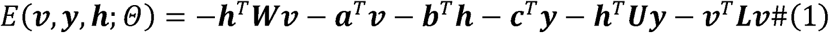

with that *Θ* = {***W, a, b, c, U, L***} represents the model parameters. Note that the first three terms in the energy function are the same in the standard RBMs in which ***W***∈ ℝ^*n*×*m*^ models the weight connections between *m* visible units and *n* hidden nodes, ***a*** ∈ ℝ^*m*^ is the bias of visible input units, and ***b*** ∈ ℝ^*n*^ is the bias of hidden nodes. The fourth and fifth terms model the contributions from the phenotype ***y*** in which ***U*** ∈ℝ^*n*×*K*^ models the weight connections between target and hidden layers, and ***c*** ∈ ℝ^*K*^ is the bias of target. The last term, a quadratic term of visible units ***v*** models two aspects contributing the energy: (1) the lateral connections among genes where ***I*** ∈ ℝ^*m*×*m*^ encodes the gene-gene relationships (i.e., adjacency matrix of a gene network) and (2) the Gaussian units of gene expression inputs. We combined these two aspects in one term because they did not affect the calculation of log-likelihood gradient in training.

The conditional probability distribution of *i*^th^ visible unit, ***v***_***i***_ with real continuous value given the hidden units and other visible units (according to lateral con^i^nections among genes), is given by:

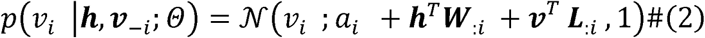

where ***L*** is a hollow matrix whose diagonal elements are all equal to zero, and ***W***_:*i*_, ***L***_:*i*_ are the *i*^th^ columns of matrices ***W*** and ***L*** respectively.

The conditional probability distribution of the output units (of binary values) given the hidden units is as follows:

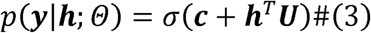

where *σ*(*x*)= (1 + *e*^-*x*^)^-l^ is the logistic sigmoid function.

The hidden units capture predictive information from both the visible inputs and the output classes. Thus, the conditional probability distribution of the hidden units (of binary values) given the visible inputs and output classes has the following form:

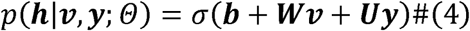

In training, because only *v* and *y* are observed, we calculated the marginal distribution represented by the model:

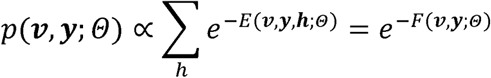

where *F*(***v, y***,; *Θ*) is the free energy function defined by *F*(***v,y***; *Θ*) = − log∑_h_ *e*^-*E*(***v,y,h***;*Θ*)^.

With the energy function aforementioned, the free energy can be further derived as follows:

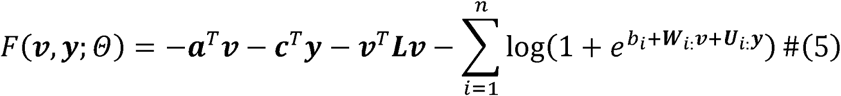

where ***W***_*i*:_ and ***U***_*i*:_ are the *i*^th^ rows of matrices ***W*** and ***L*** respectively. Furthermore, to address the “curse of dimensionality” problem (i.e., features/genes are much more than samples) in genomic datasets, we trained our classification model with an regularization. The term,, which acts on the weight connections between the visible input layer and hidden layer, and also serves as an implicit feature selection method to automatically selecting prominent genes responsible for hidden units which models the distribution over visible input units (i.e., genes). This also enables us to add more hidden units to increase the learning capacity without being overfitted. Thus, we introduced the following loss function minimization during the model training:

. The data negative log-likelihood gradient is then:
— — —
, where are generated examples from the current model’s distribution and *j* is the patient index. These generated examples can be obtained by running a Markov chain to convergence using Gibbs sampling. However, in practice, the sampling process does not wait for convergence. Instead, the samples are obtained after 5-step Gibbs sampling in our case.

#### Algorithm 1: ECMarker learning algorithm

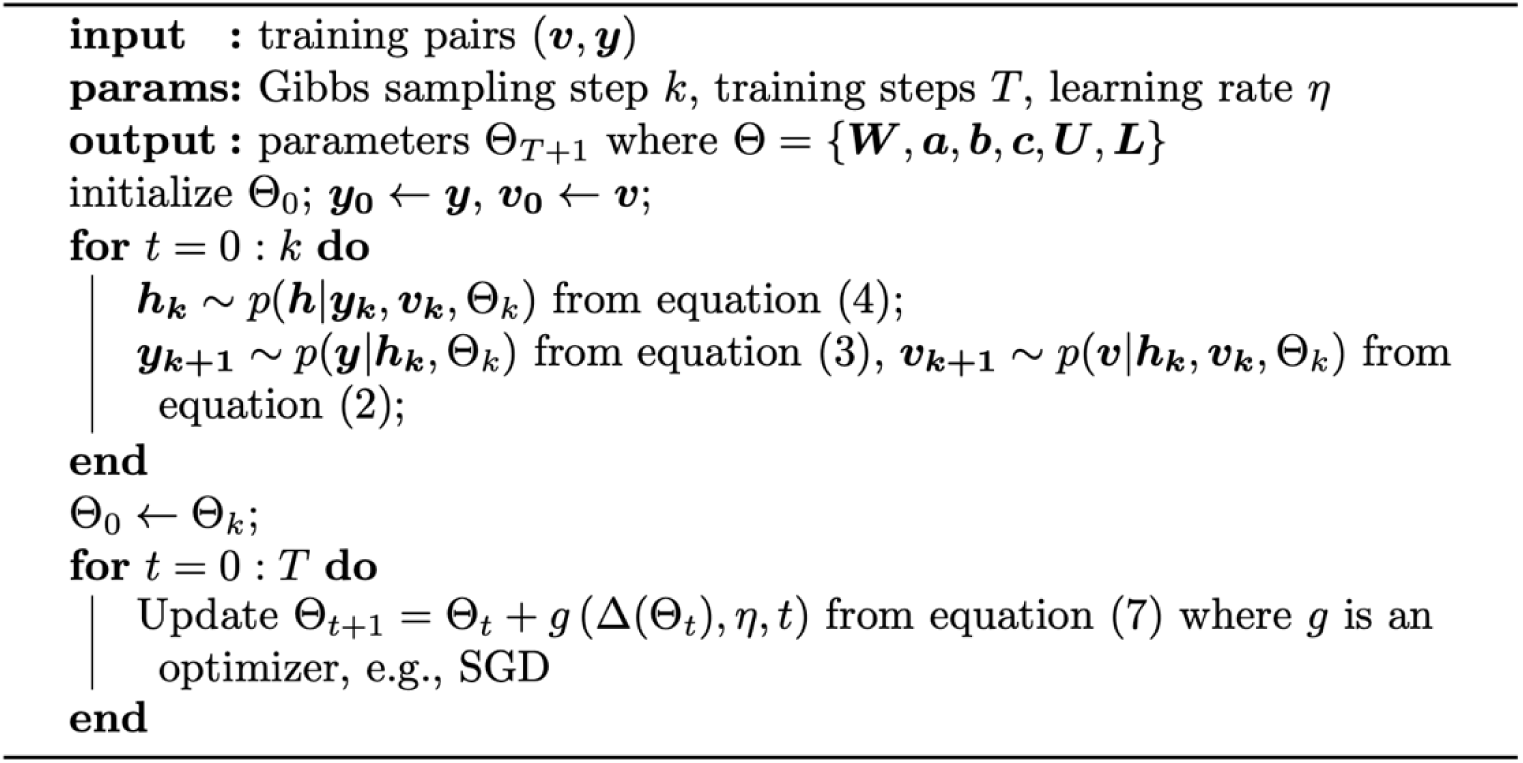

Finally, the learning procedure in the ECMarker is as follows (Algorithm 1). First, initialize, and. At the learning iteration, let be the model parameters. We generate using the Gibbs sampling. Then we update, where is the learning rate and:
—— —— —

The convergence of the algorithm toward the local optimum (since the loss function is non-convex) depends on the optimizer. We used the stochastic gradient descent (SGD) (Bottou 2010) in ECMarker, which converges to a local minimum with an explicit convergence rate of, where *T* is the number of iterations.

After training the model, the updated matrix ***L***_*t*+1_ ∈ *Θ*_*t*+1_ can be used as an adjacency matrix to construct a gene network, revealing the gene-gene relationships for predicting phenotypes (e.g., disease stages) and providing potential novel mechanistic insights.

### 2.3 Prioritization of the biomarker genes in ECMarker for phenotypes

Once the ECMarker classification model is trained, we further used a derivative-based method called integrated gradient for prioritizing input features (e.g., genes) (Sundararajan, Taly and Yan 2017). In particular, we computed the gradient of model’s prediction with respect to each individual gene to show how the output response value (i.e., early vs. late stages) changes with respect to a small change of input gene expression value. Hence, calculating these gradients for given input genes provide potential clues about which genes attribute the stage outcomes. This can be also interpreted to see which features are not selected due to *ℓ*1 regularization since the gradients for these input genes are zeros. The output response value can be computed as the posterior class probability distribution given input *v* and has the following closed form:

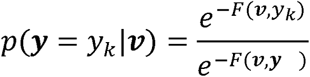

where *F*(***v, y***) is the free energy over all phenotypes in the output layer as in equation (5), and *F*(***v, y***_***k***_) is the free energy with regard to phenotype ***y*** = ***y***_*k*_ (*k* = 1, …, *K*), calculated as 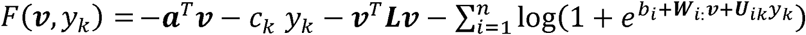. The exact gradient of this probability distribution can be calculated using the *autograd* package in PyTorch (Paszke et al. 2017). Furthermore, we define an importance score of each gene for the phenotype as the gradient of the gene to the phenotype. The higher positive scores that genes have, the more likely they contribute to predict the corresponded phenotype. Finally, given a phenotype, ECMarker prioritizes the genes for the phenotype via ranking its gene importance scores using the *Captum* package in PyTorch (Captum 2020).

### 2.4 Cancer gene expression and clinical datasets for building and testing ECMarker

We built and tested the ECMarker with the following publicly accessible gene expression datasets in lung cancer. The Gentles2015 dataset (Gentles et al. 2015) includes the log2-transformed gene expression data of 1103 non-small cell lung cancer (NSCLC) patients who had not received pre-biopsy treatment. We further imputed the missing values of the Gentles2015 dataset using the R package *impute* (Trevor Hastie 2020), and then standardized the data per sample; e.g., a mean of zero and a standard deviation of 1. Also, we grouped the patients based on their TMN stages, with (I+IA+IB) as the early stage (N = 766) and II, III, and IV as the late stages (N = 337) and divided the dataset into balanced training and testing datasets via oversampling using the R package *ROSE* (Lunardon, Menardi and Torelli 2014). The lung adenocarcinoma (LUAD) and squamous cell carcinoma (LUSC) are two of the most common subtypes of NSCLC (Herbst et al. 2018). To demonstrate the utility of ECMarker in distinct subtypes of NSCLC, we downloaded the RNA-seq gene expression datasets (FPKM values) for the LUAD and LUSC patients in TCGA (The Cancer Genome Atlas Research et al. 2013), resulting in independent validation datasets TCGA-LUAD and TCGA-LUSC. Overall, we included 741 TCGA-LUAD patients (N = 409 and N = 332 in the early and late stages, respectively) and 758 TCGA-LUSC patients (N = 380 and N = 378 in the early and late stages, respectively) with (I+IA+IB) as the early stage and the rest as the late stage. A summary on the patient numbers of various stages is in S1 Table.

### 2.5 Identification of pathways and functions associated with early and late cancer stages via enrichment analysis

We performed the enrichment analyses of the genes in the ECMarker for revealing underlying molecular mechanisms from genes to disease stages. In particular, given a phenotype (e.g., early stage), we applied the Gene Set Enrichment Analysis (GSEA) (Subramanian et al. 2005) by a R package, *fgsea* (Korotkevich, Sukhov and Sergushichev 2019) to the ranking list of all input genes based on gene importance scores for the phenotype, and found the enriched terms including pathways, functions and oncogenic signatures from all eight gene sets of the Molecular Signatures Database (MSigDB) in the GSEA (Liberzon et al. 2015, Liberzon et al. 2011). Using this enrichment analysis, we identified the enriched terms for both early and late lung cancer stages, providing mechanistic insights in lung cancer progress.

### 2.6 Survival analysis using ECMarker biomarker genes

We used the R function, *kmeans* to partition the early cancer patients into two groups using the gene expression data of top early biomarker genes in ECMarker (N=14). The survival analyses and Kaplan-Meier plots were implemented using the R package, *survival* (Therneau 2020).

### 2.7 Gene network analysis in ECMarker

The lateral connection weights (i.e., *L* matrix) from the ECMarker model how the gene-gene pairs (rather than individual genes) contribute to predict phenotypes, providing potentially additional mechanistic insights in terms of gene-gene relationships. Thus, using the *L* matrix as adjacency matrix, we further constructed a gene network from the ECMarker model for revealing potential gene regulatory relationships, especially on the transcription factors (TFs) to target genes (TGs). In addition, we compared the ECMarker gene network with the existing widely used methods such as GENIE3 (Huynh-Thu et al. 2010) that only predict gene regulatory networks (TFs to TGs) from gene expression data, without simultaneously predicting phenotypes like ECMarker. In particular, we calculated the pairwise cosine distances between same genes; i.e., a distance of *i*^th^ row vectors of ***L*** and *G* for Gene *i*, 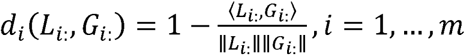, where *G* is the GENIE3’s adjacency matrix. The cosine distance ranges from 0 meaning exactly the same to 2 meaning exactly opposite. Actually, the cosine distances have been widely used to measure the similarity of vectors and matrices, revealing the structural equivalence of the same vertices (i.e., same gene) between two networks via calculating weighted common neighbors divided by the geometric mean of their degrees (Salton 1988). Furthermore, the cosine distance does not depend on the magnitude of the vector, which likely varies across different methods (e.g., ECMarker vs. GENIE3) and thus is incomparable to each other. In contrast, the other distance metrics such as the Euclidean distance or correlation take in account for the vector/matrix magnitudes, so are not chosen here for evaluating comparison. We used the Gentles2015 dataset for comparing ECMarker and GENIE3.

In addition to looking at the ECMarker network nodes and links, we also inferred the TF-TG relationships from the ECMarker network structures. In particular, we clustered the ECMarker gene network into a set of gene modules via hierarchical clustering (10 modules for the Gentles2015 dataset). The genes clustered together into a same module have strong connections for phenotype prediction, implying potential similar mechanisms such as co-regulation. Thus, we further identified the TFs with enriched target genes in each module (via TF binding sites on target gene regulatory regions) by g:Profiler (Raudvere et al. 2019) and linked them to the modular target genes. Finally, we also calculated the centralities of the ECMarker gene network using *igraph* (Csárdi and Nepusz 2006) and found the hub genes with high centrality (e.g., degree).

### 2.8 Identification of drugs targeting biomarker genes predicted by ECMarker

For discovering potential novel genome medicine using ECMarker, we identified the drugs targeting ECMarker biomarker genes of early and late cancer stages. In particular, we looked at a drug-gene database, GSCALite (Liu et al. 2018a) that has calculated and summarized the z-scores of drug-gene pairs using the method in (Rees et al. 2016) for revealing the mechanisms of action (MoA) of drugs to the target genes. The drugs with high z-scores imply potential causal mechanistic effects such as activation mechanisms and direct protein targets to the genes (Rees et al. 2016). The z-scores were the Fisher’s z-transformed correlation coefficients between gene expression and drug sensitivity (IC50 value) across all possible cancer cell lines in the GDSC database (Yang et al. 2013), and thus removed potential biased effects from specific tissue types corresponding to the cell lines. Given a phenotype (e.g., early cancer stage), we found a number of drugs for its ECMarker biomarker genes with high z-scores (FDR < 0.05), suggesting their potential effects to the phenotype (e.g., early cancer drugs).

## 3 Results

### 3.1 Lung cancer stage biomarker genes by ECMarker reveal functions on cancer development and progress

We applied ECMaker to the Gentles2015 dataset (Methods) for predicting the biomarker genes for lung cancer development and outcomes, especially for early-stage patients. In particular, we input the expression data of 10102 genes from 766 early and 766 late patients (after balancing data) in the Gentles2015 dataset. After tuning hyperparameters in this ECMarker classification model, we had: (1) the input layer containing 10102 genes; (2) the hidden layer containing 9 hidden units; (3) the output layer predicting early or late stage by a probability. Other hyperparameters were optimized as follows: train batch size = 50; learning rate = 0.1 and weight decay = 0.9 with the SGD method (Bottou 2010); *ℓ*1 penalty parameter: 0.1; number of training epoch: 1. We also performed k=10 cross-validation and found that the model has the consistent relatively high balanced accuracy values with Mean = ∼0.74 and Variance =∼ 0.001 compared to a baseline of 0.5 (for two phenotypes). Also, we applied another RBM-based model, elastic restricted Boltzmann machines (eRBMs) (Zhang et al. 2017) that does not model lateral connections at the input layer, and found that its accuracy for predicting early and late stages is just around baseline of 0.5.

After training and cross-validating models, we used the average predictive model to further calculate the gene importance scores for both early and late lung cancer stages (Methods), and prioritized the stage biomarker genes (i.e., high importance scores) in the lung cancer (Supplement File 1). As shown on Fig. 2A, a number of known lung, immunity and cancer related pathways, especially on cancer development are significantly enriched among top early biomarker genes after gene set enrichment analyses (Methods); e.g., the epithelial mesenchymal transition (EMT, *p* < 6.2e-3) (Lu and Kang 2019), the gamma delta T cell activation (*p* < 6.1e-3) (Pauza et al. 2018) and Interleukin-1 regulation (*p* < 4.0e-3) (Lewis et al. 2006). Furthermore, top early and late genes are enriched with different upregulated pathways relating to lung cancer (*p* < 0.001, Figs. 2B and 2C); e.g. lung cell differentiation and epithelium development are upregulated for the early stage, but lung cancer survival and differential markers are upregulated for the late stage. All enriched terms for early lung cancer stage are available in Supplement File 2.

**Fig. 2.**
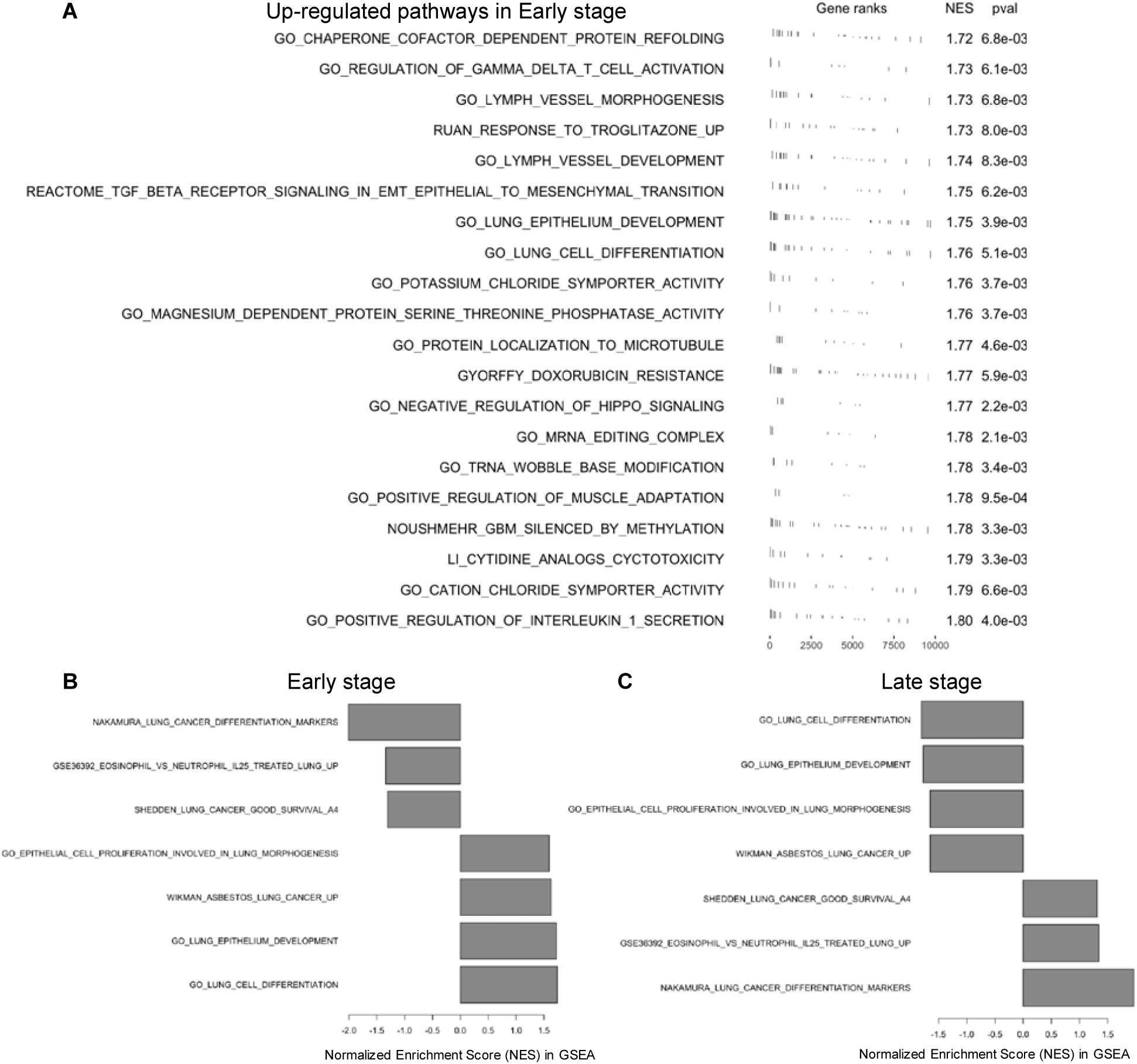
Cancer-stage biomarker genes of ECMarker reveal the biological functions and pathways associated with lung cancer and cancer development. The enrichment analyses were accomplished by the gene set enrichment analysis (GSEA) for the ranked genes by gene importance scores for early or late lung cancer stages. (**A**) The upregulated functions and pathways significantly enriched in the early stage biomarker genes with *p* < 0.01. (**B**) and (**C**) Select functions and pathways associated with lung cancer and cancer development are up- or down-regulated at different stages.

Lung cancer is also heterogeneous; e.g., non-small cell lung cancer has two major subtypes: adenocarcinoma (LUAD) and squamous cell carcinoma (LUSC) (Lucchetta et al. 2019). To test ECMarker for predicting the cancer stages in the lung cancer subtypes, we also applied ECMarker to the TCGA-LUAD and TCGA-LUSC gene expression datasets (Methods). To obtain an appropriate sample size for training ECMarker as the Gentles2015 dataset, we combined TCGA-LUAD and TCGA-LUSC together and trained one ECMarker model for classifying four phenotypes: the LUAD early and late stages and the LUSC early and late stages, aiming to reveal the specific early cancer mechanisms to lung cancer subtypes. After training and testing, the model achieved a high classification accuracy of 0.48 compared to a baseline of 0.27 (for four phenotypes). Furthermore, we found that top early-stage biomarker genes for LUAD and LUSC have significantly anti-correlated importance scores (S1 Fig.), suggesting potential distinct early cancer mechanisms across the lung cancer subtypes.

### 3.2 ECMarker biomarker genes predict clinical outcomes for early lung cancer

In addition to the genomic functions associated with cancer stages discovered by ECMarker, we also explored the relationships between stage biomarker genes and clinical outcomes of cancer patients. For example, we found that three lung cancer prognostic biomarker genes, CX3CR1, SLC15A2 and TFRC found by a recent multi-omics study (Haghjoo, Moeini and Masoudi-Nejad 2020) also have very high ECMarker importance scores for early stage (> 80% genes). In addition, we found that the early lung cancer genes, SLC15A2 and TFRC are also hub genes (degree centrality in 1% and 10%) in the gene network revealed by ECMarker. To test the capability of ECMarker for predicting clinical outcomes for early cancer, we used top early-stage biomarkers learned by ECMarker (i.e., highest importance scores for early stage, Methods) to partition the early cancer patients of Gentles2015 into two groups. We then found that two groups have significantly differential survival rates, suggesting that our ECMarker early biomarkers are able to predict early cancer survival rates (*p* < 0.004, Fig. 3A). Furthermore, we validated the top ECMarker early biomarker genes using early patients in two independent cohorts, TCGA-LUAD and TCGA-LUSC, and found that the early patients groups clustered by these biomarkers also have significantly differential survival rates (*p* < 0.0025, Figs. 3B and 3C). This demonstrates that our early biomarker genes have potential to predict survivals at the early cancer stage, suggesting the clinical interpretability of the ECMarker model.

**Fig. 3.**
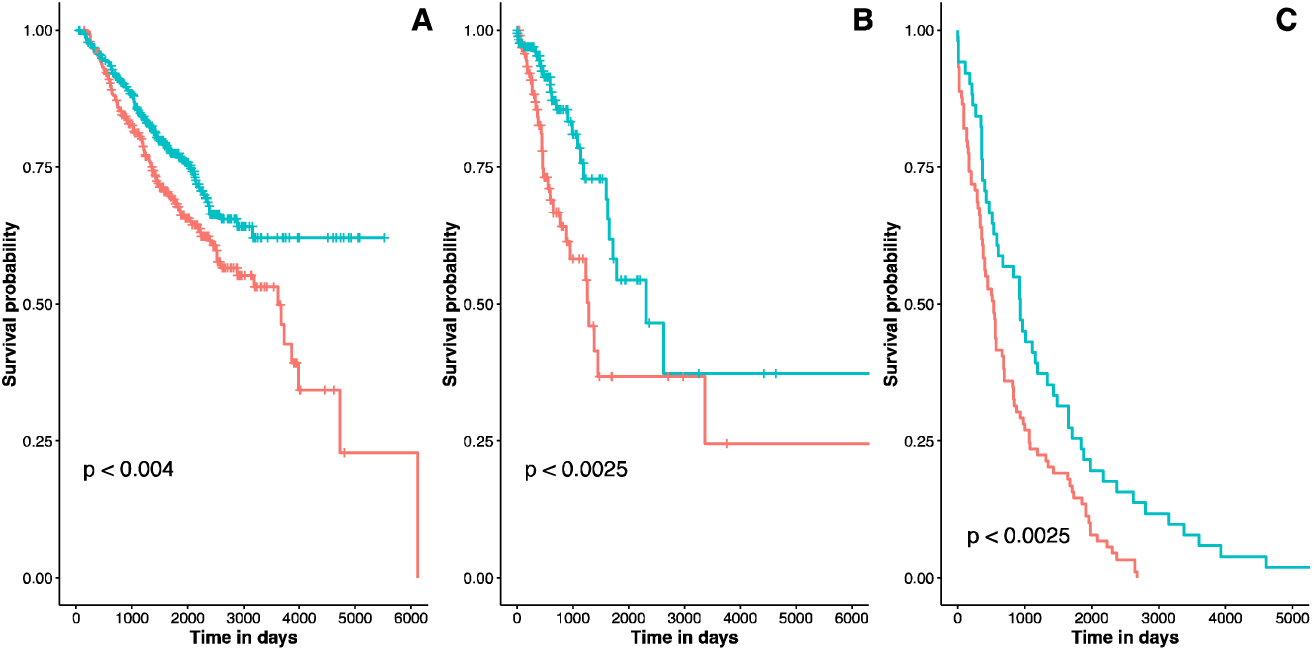
ECMarker stage biomarker genes predict early cancer survival rates. Early lung cancer patients were clustered into two groups (represented by two curves on each panel) based on the top 14 ECMarker biomarker genes for early stage lung cancer. These biomarker genes were found using the Gentles2015 dataset (Gentles et al. 2015). A Kaplan–Meier analysis showed that the early patient groups had significantly different survival rates (*p* < 0.005) as shown in Panel A. In addition, application of these biomarker genes to an independent lung cancer cohorts, TCGA-LUAD and TCGA-LUSC (The Cancer Genome Atlas Research et al. 2013), showed that the early-stage patients also had significantly different survival rates, as shown in Panel B (TCGA-LUAD) and Panel C (TCGA-LUSC) with *p* < 0.0025.

### 3.3 Gene network in the ECMarker uncovers gene regulatory mechanisms in lung cancer

In addition to individual biomarker genes, we pursued the elucidation of the molecular mechanisms that drive the functional connectivity, especially in terms of gene regulation. Gene expression, though complex for phenotypes, is controlled by gene regulatory mechanisms. In particular, various regulatory factors, such as transcription factors (TFs), control the expression of biomarker genes to coordinate cancer phenotypes so they do not behave randomly (i.e., forming a GRN). ECMarker has the capability to model gene-gene interactions so that we could extract the weight values of lateral connections to describe the relationship of any pair of genes among the input genes. A larger weight value indicated a stronger connection between genes. According to this standard, we are able to extract the learned GRN from any well-trained ECMarker classification model (Methods). Specifically using the ECMarker gene network learnt from the Gentles2015 dataset for predicting lung cancer stages, we found the subnetworks linking top early and late biomarker genes are different (Figs. 4A and 4B), suggesting potential developmental regulatory mechanisms in the lung cancer progress.

**Fig. 4.**
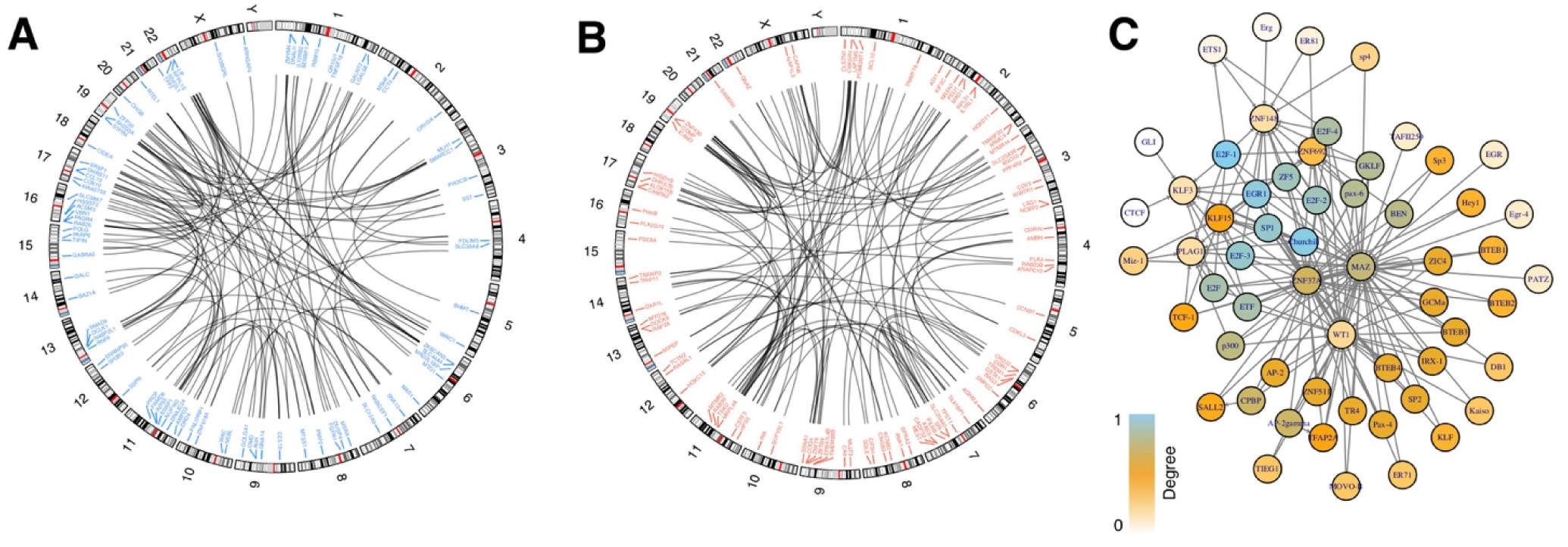
The ECMarker gene network reveals the stage-specific gene connections and regulatory networks in the lung cancer. **(A)** The ECMarker gene network for early-stage biomarker genes (gene importance score > 0.001 and top 100 links). (**B**) The ECMarker gene network for late-stage biomarker genes (gene importance score > 0.001 and top 100 links). (**C**) A regulatory network among TFs inferred from the ECMarker gene network. The nodes are TF genes. The directed edges link TFs to the target genes (which are also TFs in this case). The node color corresponds to the degree centrality of TF in the network; e.g., skyblue represents high degree, and white means low degree. Note that all TFs shown here are predicted to associated with the lung cancer development from the ECMarker model.

We found that a number of known TFs related to cancer development and oncogenes involve in the ECMarker gene network (from top 1% links), including 46 epithelial-to-mesenchymal transition (EMT) signature genes (Byers et al. 2013), lung cancer mutation genes (e.g., KRAS, BRAF, ALK, PIK3CA, AKT1, NRAS, EGFR, RET, ROS1) (Khoo et al. 2015) and the genes in the frequency of alterations and signaling pathways of the lung cancer (53 out of 83 such genes (Li et al. 2017)). Moreover, a number of gene pairs from the top ECMarker links were also previously predicted to relate to the lung cancer gene regulation. For example, there are 2676 ECMarker top pairs (117 TF, 2348 TGs) also presented in a lung cancer regulatory network predicted by the one-class support vector machine (OC-SVM) model (Zhang et al. 2018b). Also, a recent study has identified 10 oncogenic TFs and 11 tumor suppressing genes potentially required for NSCLC cell proliferation (Zhang et al. 2018a). We found that all 10 such TFs and 10 out 11 suppressing genes in our top ECMarker links (involved in 1017 pairs). In addition, a previous correlation-based analysis focusing on microRNA targets in lung cancer also predicted a set of TFs and target genes for the NSCLC (Mitra et al. 2014), and was found to have 22 TFs, 101 target genes and 13 TF-TGs presenting in the top ECMarker links. Finally, we systematically compared the lung cancer networks of ECMarker and other computational methods such as GENIE3 that only use gene expression data to predict gene regulatory networks without integrating any phenotypic information (Methods), especially on human diseases. Although ECMarker predicted a gene network specifically for predicting lung cancer development (i.e., stages), rather than generally for the lung cancer, there are still a variety of genes with high similarity between two networks (N=1369 genes, 13.6% with cosine distance < 0.35). This shows a consistency between ECMarker and GENIE3 but also implies that rest of the genes without high network similarity potentially relate to the specific developmental functions and pathways in lung cancer such as immune response, cell proliferation and differentiation (S2 Fig.).

In addition to the ECMarker network nodes and links, using the network structures (e.g., gene modules), we also identified a list of TFs and TF-TG pairs for the lung cancer and cancer development (Methods). In particular, we found a number of oncogenic TFs in our list, such as E2F genes, the cell-cycle TFs relating to tumor progression (Johnson and Schneider-Broussard 1998), and EGR genes, the TFs regulating multiple tumor suppressors (Baron et al. 2006). Furthermore, we identified a number of potential master regulators in lung cancer development using our network. For example, E2F-1, a well-known transcription factor promoting the tumor progression for many cancer types including lung cancer (Engelmann and Putzer 2012, Zhang et al. 2018b), plays a hub role (i.e., high degree) in the gene regulatory subnetwork in which target genes are TFs as well (Fig. 4C). Also, SP1, another known TF regulating lung cancer progression (Hsu et al. 2012) is in our network and also a hub gene. These hub genes in the ECMarker network imply that they regulate a number of lung cancer TFs as potential master regulators in lung cancer development. In addition, we found potential novel TFs for lung cancer development which were previously found to associate with other cancer types, such as WT1 for leukemia, kidney and prostate cancers (Hastie 2017) and MAZ, a MYC-associated zinc finger protein for pancreatic cancer (Maity et al. 2018). In addition, the target genes of some TFs are found to have significantly higher stage-specific importance scores than non-target genes (t-test *p* < 0.05), suggesting that the cancer stage associated effects of the TFs; e.g., SP1 and AP-2 for late stage, TCF-1 and ER81 for early stage.

### 3.4 ECMarker biomarker genes link to potential novel drugs for early lung cancer

We further identified a number of drugs directly affecting ECMarker stage biomarker genes, aiming to provide potential novel candidates for early cancer medicine. Using the mechanisms of actions (MoAs) of drugs to genes (Methods) (Rees et al. 2016), we identified a list of drugs for top 10 ECMarker early and late stage biomarker genes (Supplement File 3). As shown on Fig. 5, the drugs and stage biomarker genes can be in general clustered into early and late groups, suggesting the stage-specific drug effects on lung cancer development. Our analyses revealed that several known drugs for lung cancer also have high effects to our stage biomarker genes; e.g., the Type I RAF inhibitor - Dabrafenib and the Type II RAF inhibitor - AZ628 for the treatment of non-V600 BRAF mutant lung cancer (Noeparast et al. 2018) in the late stage group, and YM155 for delaying the growth of NSCLC tumor xenografts is in the early stage group (Iwasa et al. 2008).

**Fig. 5.**
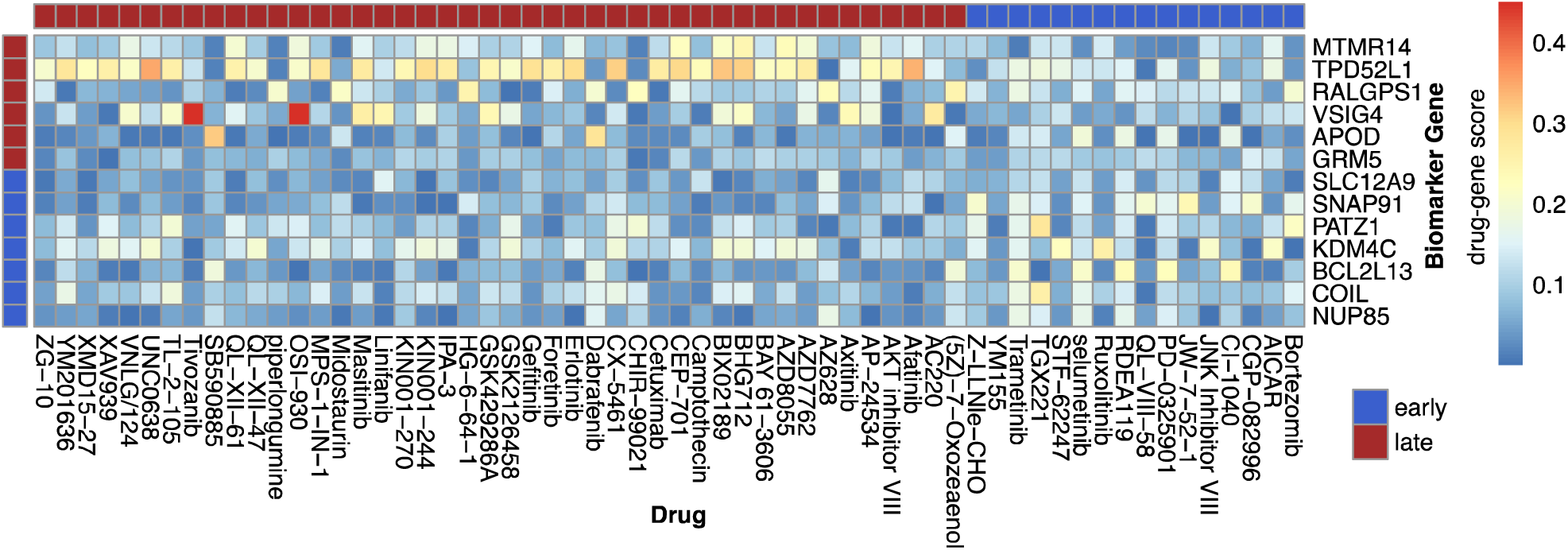
ECMarker biomarker genes discover potentially novel effective drugs for early lung cancer. The heatmap shows the effective scores of drugs to genes in terms of the mechanisms of actions (MoAs) of drugs to genes (Methods) (Rees et al. 2016). The columns are the drugs with high MoAs to at least one of top 10 ECMarker stage biomarker genes. The rows are the genes from top 10 biomarker genes targeted by the drugs (blue: early stage, red: late stage).

Furthermore, several known drugs that were not originally used for lung cancer were predicted to have significant efforts on our early-stage biomarkers; e.g., Bortezomib (Jones et al. 2010), a proteasome inhibitor ameliorating breast cancer osteolytic disease, and AICAR inhibiting the cell growth in prostate cancer cells (Digregorio et al. 2019). In addition, TAK1 inhibitor 5Z-7-oxozeaenol for the treatment of cervical cancer is found to have potential effects on our late-stage lung cancer biomarkers (Guan et al. 2017). Additionally, we observed that a few drugs previously used for multi-cancer or in clinical use for the late cancer stages are in the early stage group, suggesting their potential effects to the early lung cancer; e.g., CI-1040 and PD-0325901 for advanced non-small cell lung, breast, colon, or pancreatic cancers (Rinehart et al. 2004). Another example is Ruxolitinib, a drug used during the Phase II study in the breast cancer (Stover et al. 2018) and also possibly for NSCLC patients in all-stage to enhance oncolytic virotherapy (Patel et al. 2019). Therefore, these drugs could potentially have effects on early cancer stages.

## 4 Discussion

ECMarker is an interpretable machine learning approach, built on the SRBM and DRBM for identifying gene expression biomarkers for disease phenotypes such as cancer stages. Beyond that a variety of machine learning methods typically pursuing the high prediction accuracy from genes to phenotypes (S2 Table), we demonstrated that the ECMarker model also has biologically and clinically interpretabilities, in addition to high accuracy; e.g., it revealed the underlying regulatory mechanisms during lung cancer development and the stage biomarker genes predicted the survival rates of early cancer patients. Also, we showed that the drugs targeting the ECMarker biomarker genes are potential novel candidates for early cancer medicine. These biomarker genes comprise novel molecular candidates for early cancer diagnosis and detection, and the gene networks could potentially guide future experimental validations for early cancer mechanisms and treatments. Furthermore, ECMarker is scalable for inputting all possible genes and implicitly selecting biomarker genes for phenotypes via neural network regularization, and thus does not need any prior feature selections. Although this study applied ECMarker to lung cancer data specifically, ECMarker is a general-purpose method and can therefore be applied for other cancer types (Bailey et al. 2018) and disease types such as neurodevelopmental and neurodegenerative diseases (Li et al. 2018, De Jager et al. 2018).

This study demonstrated that we are able to build the machine learning models that are biologically interpretable. This was our first round of attempts to address the lack of interpretability and translation of machine learning applications in biology and biomedicine. Given that cancer phenotypes are driven by a variety of multi-omic mechanisms (Bailey et al. 2018), including transcriptomics, epigenomics, metabolomics, etc., multi-omic data integration and analyses for understanding cancer biology have been emerging (Rappoport and Shamir 2019). Thus, we expect to develop advanced machine learning approaches that can reveal interactions across multi-omics relating to disease phenotypes in the near future; e.g., via multiview learning approaches (Nguyen and Wang 2020). In particular, by integrating genotyping data, ECMarker can be extended to a deep hierarchical model, similar to deep neural network models (Wang et al. 2018), to predict genotype-phenotype relationships and use intermediate biological connectivity and structures inside the model to reveal possible molecular mechanisms from genotype to phenotype.

The present study focused on gene expression data at the individual tissue level. However, the cancer tissues consist of different cell types with various fitness and mutational profiles (Saadatpour et al. 2015). The continuous development of single-cell genomic and transcriptomic analyses for cancer research will enable us to explore how single cells contribute to cancer tissue expression and eventually affect phenotypes; e.g., single-cell deconvolution to estimate cell-type fractions (Baron et al. 2016, Wang et al. 2019). Integrating single-cell data into the interpretable machine learning modeling and drug association analysis might uncover novel biological mechanisms and targetable key regulators at the cellular resolution for the advancement of precision cancer medicine.

## 5 Author Contributions

D.W. conceived and designed the project. T.J., N.N. and D.W. performed data processing and analyses. T.J., N.N., F.T. and D.W. wrote the manuscript. All authors read and approved the final manuscript.

## 7 Supplementary Material

Supporting document contains Supplementary Figures 1-2 and Supplementary Tables 1-3.

Supplementary File 1 – Gene importance scores in ECMarker for early and late lung cancer stages

Supplementary File 2 - A list of the enriched functions and pathways by GSEA for early lung cancer (*p* < 0.05)

Supplementary File 3 – Drugs having potential direct effects to top 100 ECMarker stage biomarker genes in lung cancer (FDR<0.05, absolute correlation > 0.2 in GSCALite (Liu et al. 2018a))

## 8 Data Availability Statement

The derived datasets are all provided by the supplemental materials.

## Reference

Bailey, M. H., C. Tokheim, E. Porta-Pardo, S. Sengupta, D. Bertrand, A. Weerasinghe, A. Colaprico, M. C. Wendl, J. Kim, B. Reardon, P. Kwok-Shing Ng, K. J. Jeong, S. Cao, Z. Wang, J. Gao, Q. Gao, F. Wang, E. M. Liu, L. Mularoni, C. Rubio-Perez, N. Nagarajan, I. Cortes-Ciriano, D. C. Zhou, W. W. Liang, J. M. Hess, V. D. Yellapantula, D. Tamborero, A. Gonzalez-Perez, C. Suphavilai, J. Y. Ko, E. Khurana, P. J. Park, E. M. Van Allen, H. Liang, M. C. W. Group, N. Cancer Genome Atlas Research, M. S. Lawrence, A. Godzik, N. Lopez-Bigas, J. Stuart, D. Wheeler, G. Getz, K. Chen, A. J. Lazar, G. B. Mills, R. Karchin & L. Ding (2018) Comprehensive Characterization of Cancer Driver Genes and Mutations. Cell, 174, 1034–1035.

Baron, M., A. Veres, S. L. Wolock, A. L. Faust, R. Gaujoux, A. Vetere, J. H. Ryu, B. K. Wagner, S. S. Shen-Orr, A. M. Klein, D. A. Melton & I. Yanai (2016) A Single-Cell Transcriptomic Map of the Human and Mouse Pancreas Reveals Inter- and Intra-cell Population Structure. Cell Syst, 3, 346–360 e4.

Baron, V., E. D. Adamson, A. Calogero, G. Ragona & D. Mercola (2006) The transcription factor Egr1 is a direct regulator of multiple tumor suppressors including TGFbeta1, PTEN, p53, and fibronectin. Cancer Gene Ther, 13, 115–24.

Bottou, L. 2010. Large-Scale Machine Learning with Stochastic Gradient Descent. 177–186. Heidelberg: Physica-Verlag HD.

Byers, L. A., L. Diao, J. Wang, P. Saintigny, L. Girard, M. Peyton, L. Shen, Y. Fan, U. Giri, P. K. Tumula, M. B. Nilsson, J. Gudikote, H. Tran, R. J. Cardnell, D. J. Bearss, S. L. Warner, J. M. Foulks, S. B. Kanner, V. Gandhi, N. Krett, S. T. Rosen, E. S. Kim, R. S. Herbst, G. R. Blumenschein, J. J. Lee, S. M. Lippman, K. K. Ang, G. B. Mills, W. K. Hong, J. N. Weinstein, Wistuba, II, K. R. Coombes, J. D. Minna & J. V. Heymach (2013) An epithelial-mesenchymal transition gene signature predicts resistance to EGFR and PI3K inhibitors and identifies Axl as a therapeutic target for overcoming EGFR inhibitor resistance. Clin Cancer Res, 19, 279–90.

Captum. Facebook Open Source, https://captum.ai/.

Clarke, R., H. W. Ressom, A. Wang, J. Xuan, M. C. Liu, E. A. Gehan & Y. Wang (2008) The properties of high-dimensional data spaces: implications for exploring gene and protein expression data. Nat Rev Cancer, 8, 37–49.

Csárdi, G. & T. Nepusz. 2006. The igraph software package for complex network research.

De Jager, P. L., Y. Ma, C. McCabe, J. Xu, B. N. Vardarajan, D. Felsky, H. U. Klein, C. C. White, M. A. Peters, B. Lodgson, P. Nejad, A. Tang, L. M. Mangravite, L. Yu, C. Gaiteri, S. Mostafavi, J. A. Schneider & D. A. Bennett (2018) A multi-omic atlas of the human frontal cortex for aging and Alzheimer’s disease research. Sci Data, 5, 180142.

Digregorio, M., A. Lombard, P. N. Lumapat, F. Scholtes, B. Rogister & N. Coppieters (2019) Relevance of Translation Initiation in Diffuse Glioma Biology and its Therapeutic Potential. Cells, 8.

Engelmann, D. & B. M. Putzer (2012) The dark side of E2F1: in transit beyond apoptosis. Cancer Res, 72, 571–5.

Frost, J. K., W. C. Ball, Jr., M. L. Levin, M. S. Tockman, R. R. Baker, D. Carter, J. C. Eggleston, Y. S. Erozan, P. K. Gupta, N. F. Khouri & et al. (1984) Early lung cancer detection: results of the initial (prevalence) radiologic and cytologic screening in the Johns Hopkins study. Am Rev Respir Dis, 130, 549–54.

Gentles, A. J., S. V. Bratman, L. J. Lee, J. P. Harris, W. Feng, R. V. Nair, D. B. Shultz, V. S. Nair, C. D. Hoang, R. B. West, S. K. Plevritis, A. A. Alizadeh & M. Diehn (2015) Integrating Tumor and Stromal Gene Expression Signatures With Clinical Indices for Survival Stratification of Early-Stage Non–Small Cell Lung Cancer. JNCI: Journal of the National Cancer Institute, 107.

Guan, S., J. Lu, Y. Zhao, S. E. Woodfield, H. Zhang, X. Xu, Y. Yu, J. Zhao, S. Bieerkehazhi, H. Liang, J. Yang, F. Zhang & S. Sun (2017) TAK1 inhibitor 5Z-7-oxozeaenol sensitizes cervical cancer to doxorubicin-induced apoptosis. Oncotarget, 8, 33666–33675.

Haghjoo, N., A. Moeini & A. Masoudi-Nejad (2020) Introducing a panel for early detection of lung adenocarcinoma by using data integration of genomics, epigenomics, transcriptomics and proteomics. Exp Mol Pathol, 112, 104360.

Hastie, N. D. (2017) Wilms’ tumour 1 (WT1) in development, homeostasis and disease. Development, 144, 2862–2872.

Herbst, R. S., D. Morgensztern & C. Boshoff (2018) The biology and management of non-small cell lung cancer. Nature, 553, 446.

Hinton, G. E. & R. R. Salakhutdinov (2006) Reducing the dimensionality of data with neural networks. Science, 313, 504–7.

Hsu, T. I., M. C. Wang, S. Y. Chen, Y. M. Yeh, W. C. Su, W. C. Chang & J. J. Hung (2012) Sp1 expression regulates lung tumor progression. Oncogene, 31, 3973–88.

Hu, Z., J. Chen, T. Tian, X. Zhou, H. Gu, L. Xu, Y. Zeng, R. Miao, G. Jin, H. Ma, Y. Chen & H. Shen (2008) Genetic variants of miRNA sequences and non-small cell lung cancer survival. J Clin Invest, 118, 2600–8.

Huynh-Thu, V. A., A. Irrthum, L. Wehenkel & P. Geurts (2010) Inferring regulatory networks from expression data using tree-based methods. PLoS One, 5.

Iwasa, T., I. Okamoto, M. Suzuki, T. Nakahara, K. Yamanaka, E. Hatashita, Y. Yamada, M. Fukuoka, K. Ono & K. Nakagawa (2008) Radiosensitizing effect of YM155, a novel small-molecule survivin suppressant, in non-small cell lung cancer cell lines. Clin Cancer Res, 14, 6496–504.

Iyer, A. S., H. U. Osmanbeyoglu & C. S. Leslie (2017) Computational methods to dissect gene regulatory networks in cancer. Current Opinion in Systems Biology, 2, 115–122.

Jagga, Z. & D. Gupta (2014) Classification models for clear cell renal carcinoma stage progression, based on tumor RNAseq expression trained supervised machine learning algorithms. BMC proceedings, 8, S2–S2.

Johnson, D. G. & R. Schneider-Broussard (1998) Role of E2F in cell cycle control and cancer. Front Biosci, 3, d447–8.

Jones, M. D., J. C. Liu, T. K. Barthel, S. Hussain, E. Lovria, D. Cheng, J. A. Schoonmaker, S. Mulay, D. C. Ayers, M. L. Bouxsein, G. S. Stein, S. Mukherjee & J. B. Lian (2010) A proteasome inhibitor, bortezomib, inhibits breast cancer growth and reduces osteolysis by downregulating metastatic genes. Clin Cancer Res, 16, 4978–89.

Khoo, C., T. M. Rogers, A. Fellowes, A. Bell & S. Fox (2015) Molecular methods for somatic mutation testing in lung adenocarcinoma: EGFR and beyond. Transl Lung Cancer Res, 4, 126–41.

Koeffler, H. P., F. McCormick & C. Denny (1991) Molecular mechanisms of cancer. The Western journal of medicine, 155, 505–514.

Korotkevich, G., V. Sukhov & A. Sergushichev (2019) Fast gene set enrichment analysis. bioRxiv, 060012.

Larochelle, H. & Y. Bengio. 2008. Classification using discriminative restricted Boltzmann machines. In Proceedings of the 25th international conference on Machine learning, 536–543. Helsinki, Finland: Association for Computing Machinery.

Lewis, A. M., S. Varghese, H. Xu & H. R. Alexander (2006) Interleukin-1 and cancer progression: the emerging role of interleukin-1 receptor antagonist as a novel therapeutic agent in cancer treatment. J Transl Med, 4, 48.

Li, M., G. Santpere, Y. Imamura Kawasawa, O. V. Evgrafov, F. O. Gulden, S. Pochareddy, S. M. Sunkin, Z. Li, Y. Shin, Y. Zhu, A. M. M. Sousa, D. M. Werling, R. R. Kitchen, H. J. Kang, M. Pletikos, J. Choi, S. Muchnik, X. Xu, D. Wang, B. Lorente-Galdos, S. Liu, P. Giusti-Rodríguez, H. Won, C. A. de Leeuw, A. F. Pardiñas, M. Hu, F. Jin, Y. Li, M. J. Owen, M. C. O’Donovan, J. T. R. Walters, D. Posthuma, M. A. Reimers, P. Levitt, D. R. Weinberger, T. M. Hyde, J. E. Kleinman, D. H. Geschwind, M. J. Hawrylycz, M. W. State, S. J. Sanders, P. F. Sullivan, M. B. Gerstein, E. S. Lein, J. A. Knowles & N. Sestan (2018) Integrative functional genomic analysis of human brain development and neuropsychiatric risks. Science, 362, eaat7615.

Li, Z., A. A. Ivanov, R. Su, V. Gonzalez-Pecchi, Q. Qi, S. Liu, P. Webber, E. McMillan, L. Rusnak, C. Pham, X. Chen, X. Mo, B. Revennaugh, W. Zhou, A. Marcus, S. Harati, X. Chen, M. A. Johns, M. A. White, C. Moreno, L. A. Cooper, Y. Du, F. R. Khuri & H. Fu (2017) The OncoPPi network of cancer-focused protein-protein interactions to inform biological insights and therapeutic strategies. Nat Commun, 8, 14356.

Libbrecht, M. W. & W. S. Noble (2015) Machine learning applications in genetics and genomics. Nature Reviews Genetics, 16, 321.

Liberzon, A., C. Birger, H. Thorvaldsdottir, M. Ghandi, J. P. Mesirov & P. Tamayo (2015) The Molecular Signatures Database (MSigDB) hallmark gene set collection. Cell Syst, 1, 417–425.

Liberzon, A., A. Subramanian, R. Pinchback, H. Thorvaldsdottir, P. Tamayo & J. P. Mesirov (2011) Molecular signatures database (MSigDB) 3.0. Bioinformatics, 27, 1739–40.

Lindeman, N. I., P. T. Cagle, M. B. Beasley, D. A. Chitale, S. Dacic, G. Giaccone, R. B. Jenkins, D. J. Kwiatkowski, J.-S. Saldivar, J. Squire, E. Thunnissen & M. Ladanyi (2013) Molecular testing guideline for selection of lung cancer patients for EGFR and ALK tyrosine kinase inhibitors: guideline from the College of American Pathologists, International Association for the Study of Lung Cancer, and Association for Molecular Pathology. Journal of thoracic oncology : official publication of the International Association for the Study of Lung Cancer, 8, 823–859.

Liu, C. J., F. F. Hu, M. X. Xia, L. Han, Q. Zhang & A. Y. Guo (2018a) GSCALite: a web server for gene set cancer analysis. Bioinformatics, 34, 3771–3772.

Liu, J., T. Lichtenberg, K. A. Hoadley, L. M. Poisson, A. J. Lazar, A. D. Cherniack, A. J. Kovatich, C. C. Benz, D. A. Levine, A. V. Lee, L. Omberg, D. M. Wolf, C. D. Shriver, V. Thorsson, S. J. Caesar-Johnson, J. A. Demchok, I. Felau, M. Kasapi, M. L. Ferguson, C. M. Hutter, H. J. Sofia, R. Tarnuzzer, Z. Wang, L. Yang, J. C. Zenklusen, J. Zhang, S. Chudamani, J. Liu, L. Lolla, R. Naresh, T. Pihl, Q. Sun, Y. Wan, Y. Wu, J. Cho, T. DeFreitas, S. Frazer, N. Gehlenborg, G. Getz, D. I. Heiman, J. Kim, M. S. Lawrence, P. Lin, S. Meier, M. S. Noble, G. Saksena, D. Voet, H. Zhang, B. Bernard, N. Chambwe, V. Dhankani, T. Knijnenburg, R. Kramer, K. Leinonen, Y. Liu, M. Miller, S. Reynolds, I. Shmulevich, W. Zhang, R. Akbani, B. M. Broom, A. M. Hegde, Z. Ju, R. S. Kanchi, A. Korkut, J. Li, H. Liang, S. Ling, W. Liu, Y. Lu, G. B. Mills, K.-S. Ng, A. Rao, M. Ryan, J. Wang, J. N. Weinstein, J. Zhang, A. Abeshouse, J. Armenia, D. Chakravarty, W. K. Chatila, I. de Bruijn, J. Gao, B. E. Gross, Z. J. Heins, R. Kundra, K. La, M. Ladanyi, A. Luna, M. G. Nissan, A. Ochoa, S. M. Phillips, E. Reznik, F. Sanchez-Vega, C. Sander, N. Schultz, R. Sheridan, S. O. Sumer, Y. Sun, B. S. Taylor, et al. (2018b) An Integrated TCGA Pan-Cancer Clinical Data Resource to Drive High-Quality Survival Outcome Analytics. Cell, 173, 400-416.e11.

Lu, W. & Y. Kang (2019) Epithelial-Mesenchymal Plasticity in Cancer Progression and Metastasis. Dev Cell, 49, 361–374.

Lucchetta, M., I. da Piedade, M. Mounir, M. Vabistsevits, T. Terkelsen & E. Papaleo (2019) Distinct signatures of lung cancer types: aberrant mucin O-glycosylation and compromised immune response. BMC Cancer, 19, 824.

Ludwig, J. A. & J. N. Weinstein (2005) Biomarkers in Cancer Staging, Prognosis and Treatment Selection. 5, 845–856.

Lunardon, N., G. Menardi & N. Torelli (2014) ROSE: a Package for Binary Imbalanced Learning. R Journal, 6, 79–89.

Maity, G., I. Haque, A. Ghosh, G. Dhar, V. Gupta, S. Sarkar, I. Azeem, D. McGregor, A. Choudhary, D. R. Campbell, S. Kambhampati, S. K. Banerjee & S. Banerjee (2018) The MAZ transcription factor is a downstream target of the oncoprotein Cyr61/CCN1 and promotes pancreatic cancer cell invasion via CRAF-ERK signaling. J Biol Chem, 293, 4334–4349.

Mitra, R., M. D. Edmonds, J. Sun, M. Zhao, H. Yu, C. M. Eischen & Z. Zhao (2014) Reproducible combinatorial regulatory networks elucidate novel oncogenic microRNAs in non-small cell lung cancer. RNA, 20, 1356–68.

Molina, J. R., P. Yang, S. D. Cassivi, S. E. Schild & A. A. Adjei (2008) Non-Small Cell Lung Cancer: Epidemiology, Risk Factors, Treatment, and Survivorship. Mayo Clinic Proceedings, 83, 584–594.

Nguyen, N. D. & D. Wang (2020) Multiview learning for understanding functional multiomics. PLoS Comput Biol, 16, e1007677.

Noeparast, A., P. Giron, S. De Brakeleer, C. Eggermont, U. De Ridder, E. Teugels & J. De Greve (2018) Type II RAF inhibitor causes superior ERK pathway suppression compared to type I RAF inhibitor in cells expressing different BRAF mutant types recurrently found in lung cancer. Oncotarget, 9, 16110–16123.

Osindero, S. & G. Hinton. 2007. Modeling image patches with a directed hierarchy of Markov random fields. In Proceedings of the 20th International Conference on Neural Information Processing Systems, 1121–1128. Vancouver, British Columbia, Canada: Curran Associates Inc.

Paik, P. K., M. E. Arcila, M. Fara, C. S. Sima, V. A. Miller, M. G. Kris, M. Ladanyi & G. J. Riely (2011) Clinical characteristics of patients with lung adenocarcinomas harboring BRAF mutations. Journal of clinical oncology : official journal of the American Society of Clinical Oncology, 29, 2046–2051.

Pao, W., V. Miller, M. Zakowski, J. Doherty, K. Politi, I. Sarkaria, B. Singh, R. Heelan, V. Rusch, L. Fulton, E. Mardis, D. Kupfer, R. Wilson, M. Kris & H. Varmus (2004) EGF receptor gene mutations are common in lung cancers from “never smokers” and are associated with sensitivity of tumors to gefitinib and erlotinib. Proceedings of the National Academy of Sciences of the United States of America, 101, 13306–13311.

Paszke, A., S. Gross, S. Chintala, G. Chanan, E. Yang, Z. DeVito, Z. Lin, A. Desmaison, L. Antiga & A. Lerer. 2017. Automatic Differentiation in PyTorch. In NIPS 2017 Workshop on Autodiff.

Patel, M. R., A. Dash, B. A. Jacobson, Y. Ji, D. Baumann, K. Ismail & R. A. Kratzke (2019) JAK/STAT inhibition with ruxolitinib enhances oncolytic virotherapy in non-small cell lung cancer models. Cancer Gene Ther, 26, 411–418.

Pauza, C. D., M. L. Liou, T. Lahusen, L. Xiao, R. G. Lapidus, C. Cairo & H. Li (2018) Gamma Delta T Cell Therapy for Cancer: It Is Good to be Local. Front Immunol, 9, 1305.

Rahimi, A. & M. Gönen (2018) Discriminating early- and late-stage cancers using multiple kernel learning on gene sets. Bioinformatics, 34, i412–i421.

Rappoport, N. & R. Shamir (2019) Multi-omic and multi-view clustering algorithms: review and cancer benchmark. Nucleic Acids Res, 47, 1044.

Raudvere, U., L. Kolberg, I. Kuzmin, T. Arak, P. Adler, H. Peterson & J. Vilo (2019) g:Profiler: a web server for functional enrichment analysis and conversions of gene lists (2019 update). Nucleic Acids Res, 47, W191–W198.

Rees, M. G., B. Seashore-Ludlow, J. H. Cheah, D. J. Adams, E. V. Price, S. Gill, S. Javaid, M. E. Coletti, V. L. Jones, N. E. Bodycombe, C. K. Soule, B. Alexander, A. Li, P. Montgomery, J. D. Kotz, C. S. Hon, B. Munoz, T. Liefeld, V. Dancik, D. A. Haber, C. B. Clish, J. A. Bittker, M. Palmer, B. K. Wagner, P. A. Clemons, A. F. Shamji & S. L. Schreiber (2016) Correlating chemical sensitivity and basal gene expression reveals mechanism of action. Nat Chem Biol, 12, 109–16.

Rinehart, J., A. A. Adjei, P. M. Lorusso, D. Waterhouse, J. R. Hecht, R. B. Natale, O. Hamid, M. Varterasian, P. Asbury, E. P. Kaldjian, S. Gulyas, D. Y. Mitchell, R. Herrera, J. S. Sebolt-Leopold & M. B. Meyer (2004) Multicenter phase II study of the oral MEK inhibitor, CI-1040, in patients with advanced non-small-cell lung, breast, colon, and pancreatic cancer. J Clin Oncol, 22, 4456–62.

Saadatpour, A., S. Lai, G. Guo & G. C. Yuan (2015) Single-Cell Analysis in Cancer Genomics. Trends Genet, 31, 576–586.

Salton, G. 1988. Automatic text processing : the transformation, analysis, and retrieval of information by computer. Reading, Mass.: Addison-Wesley.

Siegel, R. L., K. D. Miller & A. Jemal (2018) Cancer statistics, 2018. CA Cancer J Clin, 68, 7–30.

Statnikov, A., L. Wang & C. F. Aliferis (2008) A comprehensive comparison of random forests and support vector machines for microarray-based cancer classification. BMC Bioinformatics, 9, 319.

Stover, D. G., C. R. Gil Del Alcazar, J. Brock, H. Guo, B. Overmoyer, J. Balko, Q. Xu, A. Bardia, S. M. Tolaney, R. Gelman, M. Lloyd, Y. Wang, Y. Xu, F. Michor, V. Wang, E. P. Winer, K. Polyak & N. U. Lin (2018) Phase II study of ruxolitinib, a selective JAK1/2 inhibitor, in patients with metastatic triple-negative breast cancer. NPJ Breast Cancer, 4, 10.

Subramanian, A., P. Tamayo, V. K. Mootha, S. Mukherjee, B. L. Ebert, M. A. Gillette, A. Paulovich, S. L. Pomeroy, T. R. Golub, E. S. Lander & J. P. Mesirov (2005) Gene set enrichment analysis: A knowledge-based approach for interpreting genome-wide expression profiles. Proceedings of the National Academy of Sciences, 102, 15545–15550.

Sundararajan, M., A. Taly & Q. Yan. 2017. Axiomatic attribution for deep networks. In Proceedings of the 34th International Conference on Machine Learning - Volume 70, 3319–3328. Sydney, NSW, Australia: JMLR.org.

The Cancer Genome Atlas Research, N., K. Chang, C. J. Creighton, C. Davis, L. Donehower, J. Drummond, D. Wheeler, A. Ally, M. Balasundaram, I. Birol, Y. S. N. Butterfield, A. Chu, E. Chuah, H.-J. E. Chun, N. Dhalla, R. Guin, M. Hirst, C. Hirst, R. A. Holt, S. J. M. Jones, D. Lee, H. I. Li, M. A. Marra, M. Mayo, R. A. Moore, A. J. Mungall, A. G. Robertson, J. E. Schein, P. Sipahimalani, A. Tam, N. Thiessen, R. J. Varhol, R. Beroukhim, A. S. Bhatt, A. N. Brooks, A. D. Cherniack, S. S. Freeman, S. B. Gabriel, E. Helman, J. Jung, M. Meyerson, A. I. Ojesina, C. S. Pedamallu, G. Saksena, S. E. Schumacher, B. Tabak, T. Zack, E. S. Lander, C. A. Bristow, A. Hadjipanayis, P. Haseley, R. Kucherlapati, S. Lee, E. Lee, L. J. Luquette, H. S. Mahadeshwar, A. Pantazi, M. Parfenov, P. J. Park, A. Protopopov, X. Ren, N. Santoso, J. Seidman, S. Seth, X. Song, J. Tang, R. Xi, A. W. Xu, L. Yang, D. Zeng, J. T. Auman, S. Balu, E. Buda, C. Fan, K. A. Hoadley, C. D. Jones, S. Meng, P. A. Mieczkowski, J. S. Parker, C. M. Perou, J. Roach, Y. Shi, G. O. Silva, D. Tan, U. Veluvolu, S. Waring, M. D. Wilkerson, J. Wu, W. Zhao, T. Bodenheimer, D. N. Hayes, A. P. Hoyle, S. R. Jeffreys, L. E. Mose, J. V. Simons, M. G. Soloway, S. B. Baylin, B. P. Berman, M. S. Bootwalla, L. Danilova, et al. (2013) The Cancer Genome Atlas Pan-Cancer analysis project. Nature Genetics, 45, 1113.

A Package for Survival Analysis in R. CRAN. impute: impute: Imputation for microarray data. R package version 1.62.0.

Wang, D., S. Liu, J. Warrell, H. Won, X. Shi, F. C. P. Navarro, D. Clarke, M. Gu, P. Emani, Y. T. Yang, M. Xu, M. J. Gandal, S. Lou, J. Zhang, J. J. Park, C. Yan, S. K. Rhie, K. Manakongtreecheep, H. Zhou, A. Nathan, M. Peters, E. Mattei, D. Fitzgerald, T. Brunetti, J. Moore, Y. Jiang, K. Girdhar, G. E. Hoffman, S. Kalayci, Z. H. Gümüş, G. E. Crawford, P. Roussos, S. Akbarian, A. E. Jaffe, K. P. White, Z. Weng, N. Sestan, D. H. Geschwind, J. A. Knowles & M. B. Gerstein (2018) Comprehensive functional genomic resource and integrative model for the human brain. Science, 362, eaat8464.

Wang, X., J. Park, K. Susztak, N. R. Zhang & M. Li (2019) Bulk tissue cell type deconvolution with multi-subject single-cell expression reference. Nat Commun, 10, 380.

Xiao, Y., J. Wu, Z. Lin & X. Zhao (2018) A deep learning-based multi-model ensemble method for cancer prediction. Comput Methods Programs Biomed, 153, 1–9.

Yang, W., J. Soares, P. Greninger, E. J. Edelman, H. Lightfoot, S. Forbes, N. Bindal, D. Beare, J. A. Smith, I. R. Thompson, S. Ramaswamy, P. A. Futreal, D. A. Haber, M. R. Stratton, C. Benes, U. McDermott & M. J. Garnett (2013) Genomics of Drug Sensitivity in Cancer (GDSC): a resource for therapeutic biomarker discovery in cancer cells. Nucleic Acids Res, 41, D955–61.

Zhang, D. L., L. W. Qu, L. Ma, Y. C. Zhou, G. Z. Wang, X. C. Zhao, C. Zhang, Y. F. Zhang, M. Wang, M. Y. Zhang, H. Yu, B. B. Sun, S. H. Gao, X. Cheng, M. Z. Guo, Y. C. Huang & G. B. Zhou (2018a) Genome-wide identification of transcription factors that are critical to non-small cell lung cancer. Cancer Lett, 434, 132–143.

Zhang, S., M. Li, H. Ji & Z. Fang (2018b) Landscape of transcriptional deregulation in lung cancer. BMC Genomics, 19, 435.

Zhang, S., M. Liang, Z. Zhou, C. Zhang, N. Chen, T. Chen & J. Zeng (2017) Elastic restricted Boltzmann machines for cancer data analysis. Quantitative Biology, 5, 159–172.

